# Superior dispersal ability leads to persistent ecological dominance by *Candida pseudoglaebosa* in the *Sarracenia purpurea* metacommunity

**DOI:** 10.1101/387753

**Authors:** Primrose J. Boynton, Celeste N. Peterson, Anne Pringle

## Abstract

A large number of descriptive surveys document changes in microbial communities over time, but direct evidence for the ecological processes mediating succession or causing ecological dominance remains rare. Differential dispersal may be a key mechanism. We surveyed microbial diversity within a metacommunity of pitchers of the model carnivorous plant *Sarracenia purpurea* and discovered the yeast *Candida pseudoglaebosa* as ecologically dominant. Its frequency in the metacommunity increased over the growing season, and it was not replaced by other taxa. We next measured its competitive ability in a manipulative laboratory experiment and tracked its dispersal over time in nature. Despite its dominance, *C. pseudoglaebosa* is not a superior competitor. Instead, it is a superior disperser: it arrives in pitchers earlier, and disperses into more pitchers, than other taxa. Differential dispersal across the spatially structured metacommunity of individual pitchers emerges as a key driver of the continuous dominance of *C. pseudoglaebosa* during succession.

## Introduction

Primary microbial succession occurs when a microbial community colonizes and develops on a newly available substrate (Fierer *et al*. 2010). The advent of high-throughput sequencing has revolutionized observational studies of microbial succession, enabling researchers to describe the development of microbial communities in fine detail (Copeland *et al.* 2015, Boynton & Greig 2016, Koenig *et al.* 2011, Cutler *et al*. 2014). A variety of successional patterns have been observed. For example, taxon diversity can increase, decrease, or randomly vary with successional time (Cutler *et al.* 2014, Zumsteg *et al.* 2012, Redford & Fierer 2009). However, researchers commonly observe replacement of early-successional taxa by late-successional taxa (*e.g.,* Boynton & Greig 2016, Wolfe *et al.* 2014). An ongoing challenge in microbial ecology is to connect observed ecological patterns to the ecological processes responsible for the patterns.

The development of ecological dominance by one or a few microbes over successional time is a particularly intriguing phenomenon (Boynton & Greig, 2016, Findley *et al.* 2013, Jiao *et al.* 2016), in part because of its obvious parallels to plant and animal ecology (Hillebrand *et al.* 2008). Ecological dominance is apparent when one or a few species comprise most of the individuals or biomass in a community (Hillebrand *et al.* 2008). Environmental filtering, superior competitive ability of the dominant species, and ecosystem engineering can all cause dominance (Wolfe *et al.* 2014, Albergaria *et al*. 2010, Goddard 2008, Nissen & Arneborg 2003, Williams *et al*. 2015). However, the dynamics of microbial dominance have been primarily studied in domesticated systems, and the most frequently reported mechanisms may not be responsible for dominance in all, or even most, natural microbial communities. For example, environmental filtering and competition may be more important in systems where human beings have designed environments to favor domesticated microbes (Wolfe & Dutton 2015) and less important in natural environments with heterogeneous environmental conditions.

An overlooked mechanism driving ecological dominance in natural systems may be dispersal. When primary succession occurs on a sterile substrate, all members of the microbial community must first disperse onto the new substrate before establishing in the community. A good disperser may prevent the establishment of other community members through priority effects if it arrives in a habitat first, either by pre-empting or modifying available niches (Fukami 2015). Additionally, good dispersers in nearby communities can impact a microbial community during succession by producing propagules that then disperse into the developing community (Wilson & McTammany 2016).

Dispersal may emerge as a key driver of ecological dominance in microbial metacommunities. Metacommunities are physically structured groups of communities in which individual communities are spatially isolated from one another but linked through dispersal (Logue *et al.* 2011, Holyoak *et al.* 2005). Community assembly in metacommunities is a function of dispersal among communities and ecological processes occurring inside each community. Ecological theory explains how dispersal can interact with intracommunity processes to maintain metacommunity diversity (Leibold *et al.* 2004, Holyoak *et al.* 2005). For example, populations occupying low-quality environmental patches can be maintained by dispersal from high-quality patches (“mass effects”), or fitness trade-offs between competitive ability and dispersal ability can mediate species diversity (“patch dynamics”). Theory predicts that dispersal and competition interact during succession in the individual component communities of a metacommunity and the result is a hump-shaped relationship between successional time and species richness (Moquet *et al.* 2003, Sferra *et al.* 2017). Species richness is predicted to be low early in a component community’s age, to increase as more species disperse into the community, and then to decrease as competitive interactions remove species from the community. Although rarely documented, it is also possible that a particularly good disperser will become dominant and remain dominant (and therefore decreases community diversity) in a metacommunity solely as a result of its dispersal ability.

We investigated the contribution of dispersal to ecological dominance over the course of natural microbial succession in pitchers of the carnivorous pitcher plant *Sarracenia purpurea*. *S. purpurea* is a perennial plant native to bogs and savannas in northern and eastern North America (Buckley *et al.* 2003). Each *S. purpurea* plant produces modified vase-shaped leaves, or pitchers (Figure 1a), annually. At first, developing pitchers are entirely closed, sterile chambers (Peterson *et al.* 2008). Once mature, the top portion of each pitcher opens, and open pitchers accumulate rainwater to form small pools of water (phytotelmata). Potential prey (ants and other small insects) are attracted to pitchers (Bennett & Ellison 2009); some prey fall into pitchers and drown, and are then shredded, decomposed, and mineralized by a food web of microorganisms and invertebrates (reviewed in Ellison *et al.* 2003). The pitcher microbial community includes bacteria, algae, and fungi, including culturable yeasts (Cochran-Stafira & von Ende 1998). Pitchers are individual communities within a metacommunity of other pitchers on the same and on different plants.

**Figure 1:**
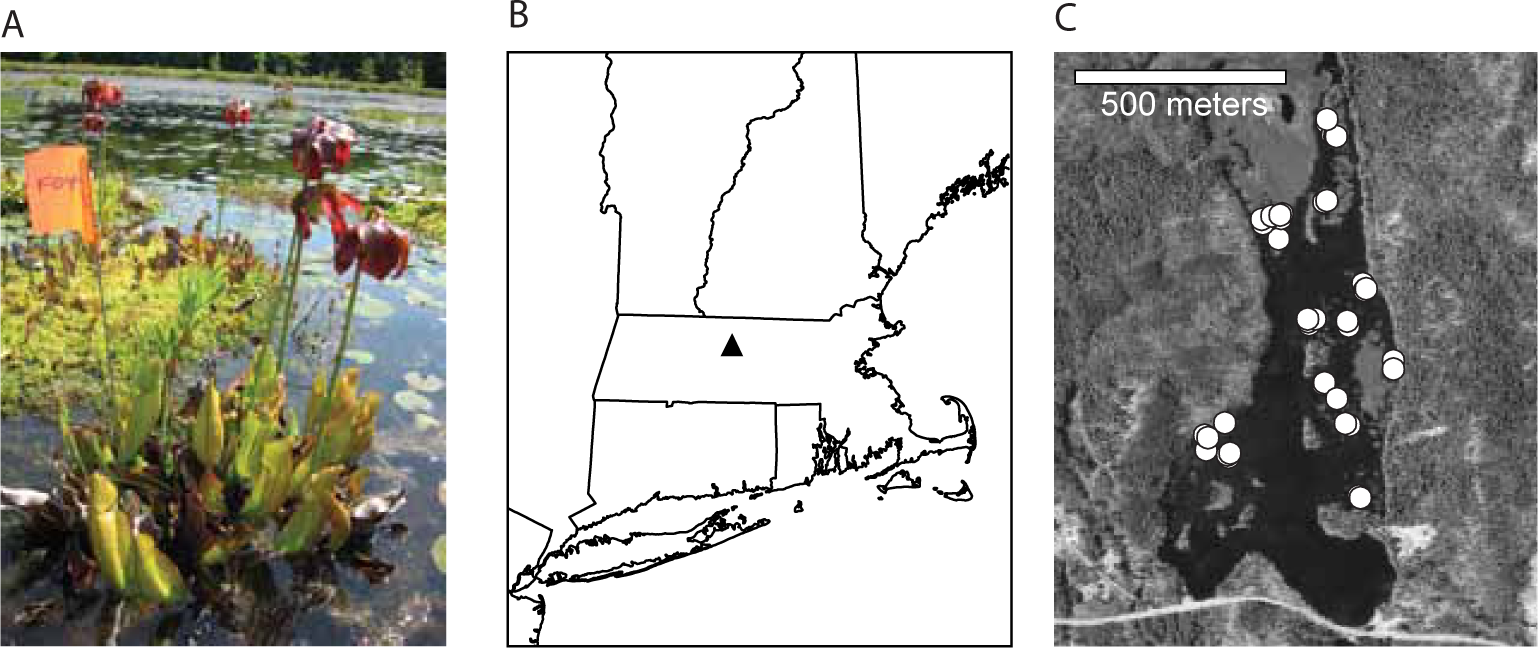
Example *S. purpurea* plant and study location. (A) One of the study pitcher plants at the edge of a *Sphagnum* island. This photograph was taken early in the growth season and both opened and unopened pitchers are visible. (B) The location of Harvard Pond in Massachusetts, USA. (C) Locations of the 43 pitchers sampled for this study. Each white circle represents one pitcher. Note that some pitchers were close enough to one another that the white circles overlap. Maps were created using ArcMap™ Version 9.2 (ESRI 2006); map data are from the Office of Geographic Information, Commonwealth of Massachusetts Information Technology Division (2000) and the National Atlas of the United States (2006).

We followed fungal succession within individual pitchers in a *Sphagnum* bog in central Massachusetts, where the pitcher growing season lasts for about three months (although pitchers persist for longer and can overwinter, Judd 1959). We first documented that a single yeast species, *Candida pseudoglaebosa*, was numerically dominant in the metacommunity throughout the growing season, and its frequency increased between early and later-successional stages of individual pitchers. We next investigated the ecological processes leading to *C. pseudoglaebosa* dominance. Unlike dominant yeasts in many other systems, *C. pseudoglaebosa* was not an especially good competitor against other tested yeasts. However, it was an especially good disperser. It is one of the first fungi to arrive in pitchers and it maintained high frequency in the pitcher plant metacommunity over the course of the season, even as its frequency within individual pitchers ranged from completely absent to over 90% of sequences. The apparently superior dispersal ability of *C. pseudoglaebosa* leads it to dominate the pitcher fungal metacommunity, demonstrating that dispersal ability, like competitive ability, is an important contributor to ecological dominance in microbial communities.

## Results

### Fungal communities in pitchers change over successional time

To understand how fungal communities develop in pitchers, we first sequenced entire fungal communities in seventeen pitchers over the course of a growing season. Before selecting pitchers to track for sequencing, we identified 43 unopened pitchers on *Sphagnum* islands in Harvard Pond, located in Petersham, Massachusetts (Figure 1). We recorded the opening date of each pitcher and collected water from each pitcher at four days, seven days (“one week”), 34-42 days (“one month”), and 66-74 days (“two months”) after opening. At the two month time point, insect herbivores, including moth larvae (Atwater *et al.* 2006), had destroyed ten of the original 43 pitchers, and we could only sample water from 33 pitchers. In the sampled pitchers, the presence of fungal DNA was assayed using the ITS1F/ITS4 primer pair (Gardes & Bruns 1993, White *et al.* 1990) Fungi were detectable starting from the first measured timepoint at four days (in 33% of sampled plants), and were widespread after one week, one month, and two months (in 91%, 95%, and 73% of sampled plants, respectively). Seventeen of the pitchers contained detectable fungal DNA at every time point from one week to two months. We sequenced fungal DNA from these seventeen pitchers at all available time points, including four days if available, using 454 sequencing of PCR amplicons of the internal transcribed spacer (ITS) ribosomal region.

Fungal succession varied among pitchers. While community composition changed significantly with time (Figure 2A), only a small amount of variation in community composition was due to variation in time (distance-based redundancy analysis adjusted R^2^ = 0.03, F = 1.96, df = 1,27, p = 0.011). This small influence of time on community composition was likely a result of high variation among pitchers. Succession followed two trajectories—five pitchers (hereafter referred to as “pitcher group 1”) followed a different successional trajectory from the other twelve (“pitcher group 2”)—and there was considerable variation among communities within each trajectory (Figure 2B). Distance among pitchers did not explain significant variation in community composition (partial mantel test of community composition on space controlling for time, mantel statistic r = −0.04, significance = 0.715).

**Figure 2:**
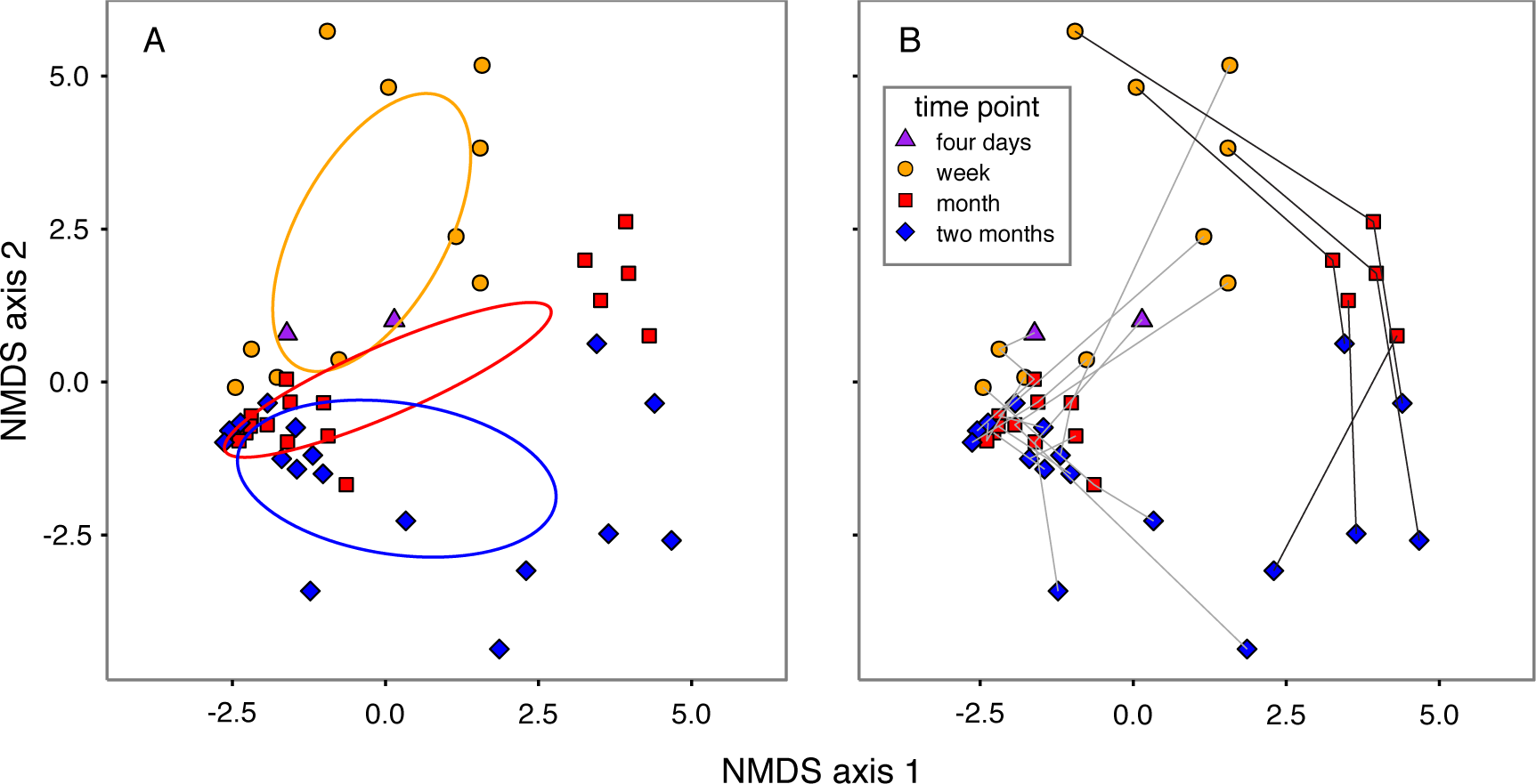
Non-metric Multidimensional Scaling (NMDS) plots of pitcher plant community similarities. Similarities of community OTU compositions were calculated using the Jaccard metric (Jaccard 1912). Purple triangles represent four-day-old communities, orange circles represent week-old communities, red squares represent approximately one-month old communities, and blue diamonds represent approximately two-month-old communities. (A) NMDS plot with similarities among time points highlighted. Ellipses depict 95% confidence intervals of the centroid of each time point. No ellipse is depicted for four-day-old communities because only two were measured. (B) NMDS plot with individual pitchers highlighted. Lines connect measurements for each pitcher. Fungal communities in pitcher group 1 are represented with black lines and fungal communities in pitcher group 2 are represented with gray lines. All points are located at the same coordinates in (A) and (B).

Despite the observed variation in fungal community composition, diversity decreased, on average, in pitchers between four days and two months (Figure 3). To determine a sample’s diversity, we calculated Hill numbers of orders 0 to 2 (^0^D to ^2^D) for each sample after rarefaction to 1140 sequences per sample. Hill numbers of different orders give community diversity with an emphasis on rare species (low orders) or common species (high orders) (Hill 1973, Chao *et al.* 2014). ^0^D, ^1^D, and ^2^D are equal to operational taxonomic unit (OTU) richness, the exponent of Shannon diversity, and the inverse of Simpson’s index, respectively. Diversity as indicated by all calculated Hill numbers decreased between early and late timepoints: on average, ^0^D declined significantly from 42.5 within the first week (including four day and one week timepoints) to 23.1 after two months (t=-3.6, df=27, p=0.001); ^1^D declined from 14.2 to 5.2 (t=-3.8, df=27, p=0.0008); and ^2^D declined from 8.9 to 3.2 (t=3.3, df=27, p=0.003).

**Figure 3:**
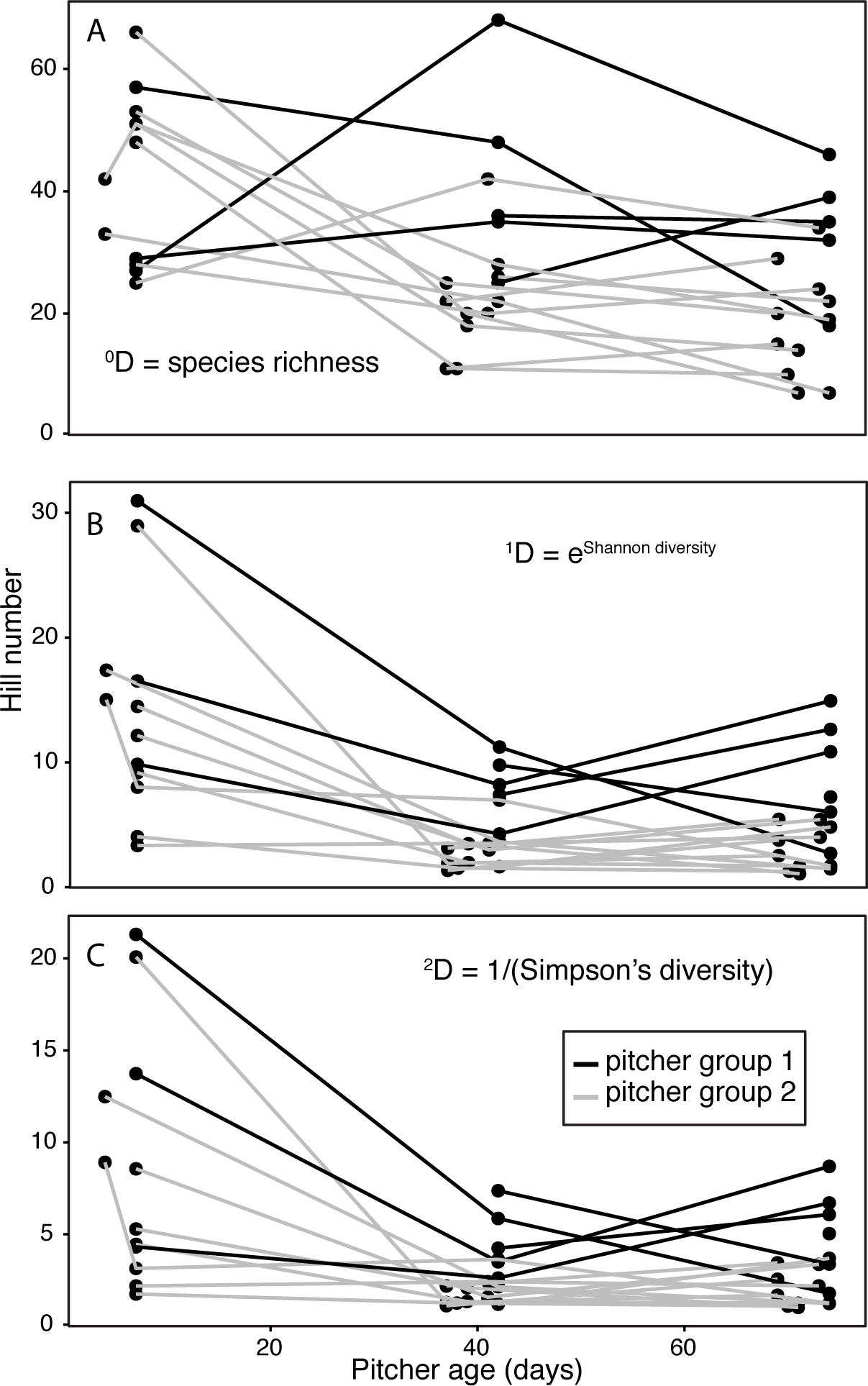
Hill numbers of orders A) 0, B) 1, and C) 2 in pitchers over time. Data points for communities in the same pitcher are connected with lines. Black lines connect points for pitcher group 1 and gray lines connect points for pitcher group 2.

### C. pseudoglaebosa is the dominant fungal taxon in pitchers throughout succession

*C. pseudoglaebosa*, in the class Saccharomycetes, was the numerically dominant taxon in the metacommunity, but was not dominant in every pitcher (Figure 4). In the metacommunity, *C. pseudoglaebosa* was more frequent than any other taxon at every time point and its frequency increased between early- and late-successional timepoints. Its metacommunity frequency increased from 22% of total sequences at four days to 58% at two months (Figure 4A). However, its frequency did not increase over time in every pitcher. Depending on the pitcher, *C. pseudoglaebosa*’s within-pitcher frequency increased over the season, decreased over the season, peaked midway through the season, or dipped midway through the season (Figure 4B). We cannot say whether these increases or decreases in *C. pseudoglaebosa* sequence frequency reflect changes in the total cell numbers because we did not measure cell numbers or fungal biomass in pitchers. Pitcher group 1 never contained appreciable *C. pseudoglaebosa*: each pitcher in group 1 contained less than 1.7% *C. pseudoglaebosa* sequences regardless of the sampled time point (Figure 4B). C. pseudoglaebosa *is not a superior competitor, but has complex interactions with other yeasts*

**Figure 4:**
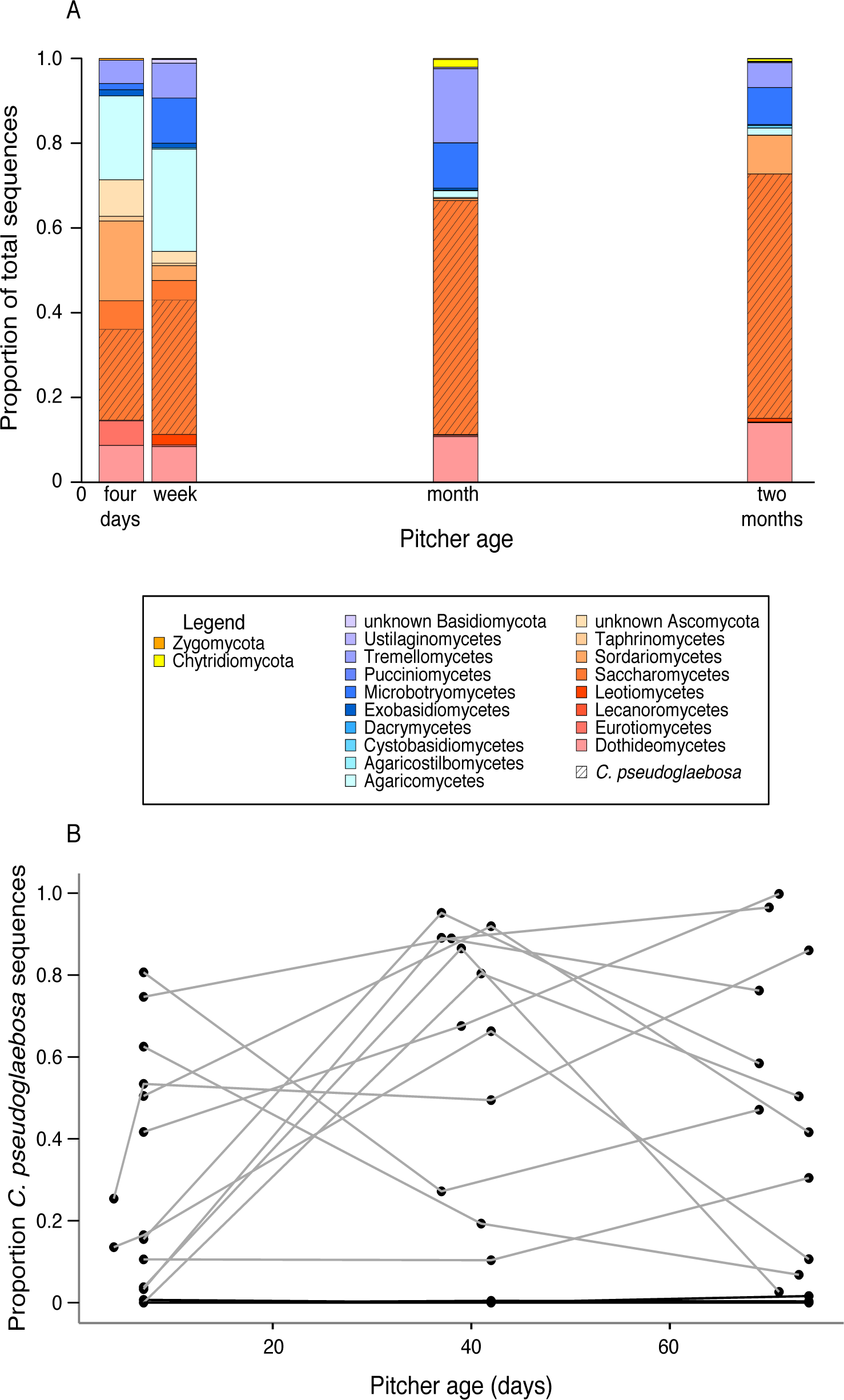
Taxon diversity in pitchers over time. Proportions are reported based on non-rarefied OTU assignments. (A) Taxon diversity in the entire bog metacommunity. Colored bars represent proportions of total sequences for each fungal class (or phylum for basal fungal lineages). The hatched area represents total *C. pseudoglaebosa* frequency for each time point. Note that *C. pseudoglaebosa* is in the class Saccharomycetes and represents over 99% of Saccharomycetes sequences at the one and two month time points. (B) *C. pseudoglaebosa* sequence frequency in individual pitcher communities. Data points for communities in the same pitcher are connected with lines. Black lines connect points for pitcher group 1 and gray lines connect points for pitcher group 2. Data for this figure is non-rarefied OTU data; for rarefied data, see Figure 4 Figure Supplement 1.

To better understand how interactions with other yeasts might influence the frequency of *C. pseudoglaebosa*, we grew *C. pseudoglaebosa* and other potentially interacting pitcher yeasts in laboratory microcosms. We followed the strategy advocated by Goldberg and Werner (1983), who suggested determining interacting species’ effects on one another by measuring organism performance as the number of interacting individuals increases. We inoculated microcosms with all possible pairs of three culturable pitcher yeasts (*C. pseudoglaebosa*, *Rhodotorula babjevae*, and *Moesziomyces aphidis*). *C. pseudoglaebosa* represented 41%, *R. babjevae* represented 2%, and *M. aphidis* represented 0.06% of total sequences in the sequencing dataset. Each low nutrient microcosm contained a focal yeast, which was inoculated as a fixed number of cells, and an interactor, which was inoculated as a varying number of cells. We then let the pairs of yeasts grow in the microcosms and investigated the effects of interactors on each focal yeast using regressions. We evaluated interaction qualities based on the direction (increasing or decreasing focal yeast yield with more interacting cells) and shape (linear or polynomial) of each regression, and we evaluated differences between interactor yeasts based on whether adding interactor yeast identity to each regression improved its fit.

Interactions between pitcher plant yeasts ranged from facilitation to competition, depending on the identities of the yeasts and the number of interactor cells present. Under microcosm conditions, interactions between focal yeasts and interactors were polynomial when the focal yeast was *C. pseudoglaebosa* or *M. aphidis* (Figures 5A, B, Figure 5 figure supplemental tables 1-4): both yeasts were facilitated by small numbers of co-inoculated cells, but their growth was impeded by larger numbers of co-inoculated cells. Note that we detected facilitation of *C. pseudoglaebosa* by *R. babjevae* when few *R. babjevae* cells were inoculated, but we did not inoculate *M. aphidis* in small enough numbers to confirm *M. aphidis* facilitation of *C. pseudoglaebosa* (Figure 5A). At high numbers of co-inoculated cells, *M. aphidis* had a more detrimental impact on *C. pseudoglaebosa* than *R. babjevae* (F=6.79, df=1,50, p = 0.012, Figure 5, figure supplemental table 2). In contrast, the two interactors of *M. aphidis* had similar effects on its yield: at low and intermediate inoculum sizes, both *R. babjevae* and *C. pseudoglaebosa* promoted *M. aphidis* growth, but at high inoculum sizes, both interactors inhibited *M. aphidis* growth (Figure 5B). Interactions between *R. babjevae* and interactor yeasts were linear (Figure 5C, figure supplemental tables 5-6): *R. babjevae* yield was impeded by interactors regardless of the number of interactor cells present, and *C. pseudoglaebosa* had a more detrimental impact on *R. babjevae* than *M. aphidis* did (F=86.07, df=1,58, p < 0.001, Figure 5, figure supplemental table 6).

**Table 1:**
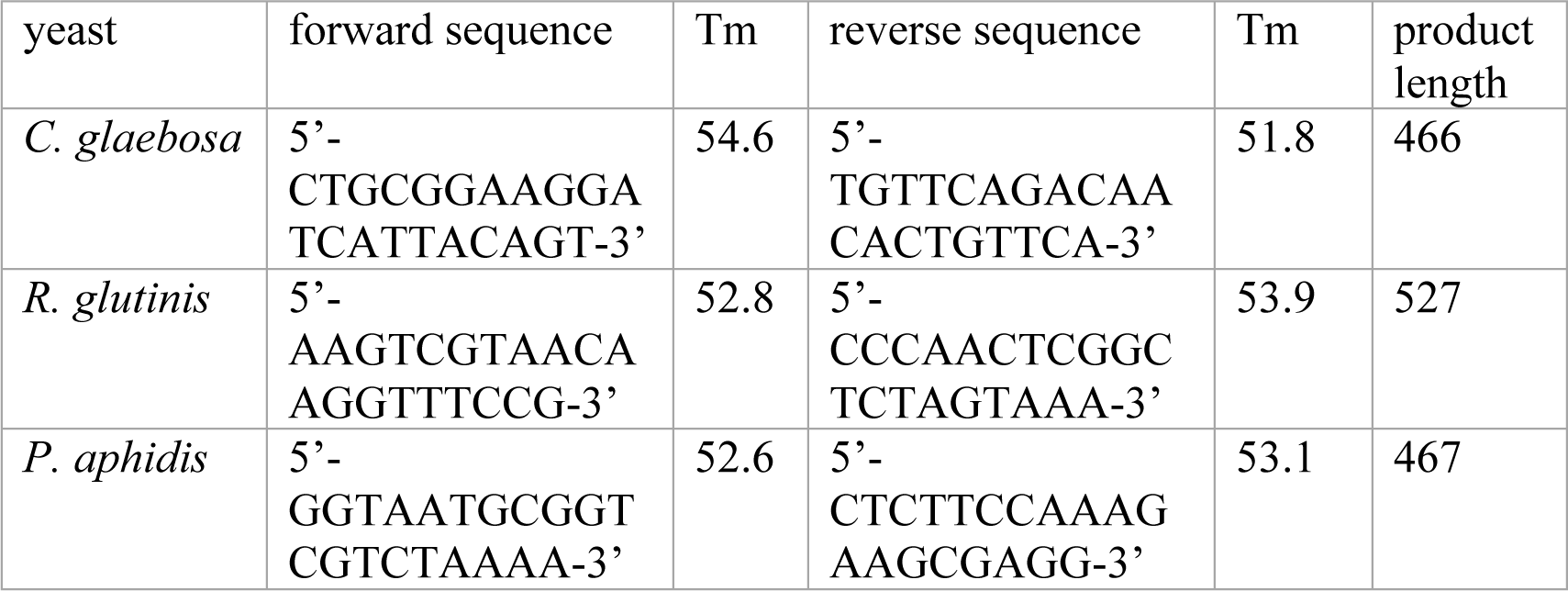
Taxon-specific PCR primer sequences used to detect pitcher yeasts

**Figure 5:**
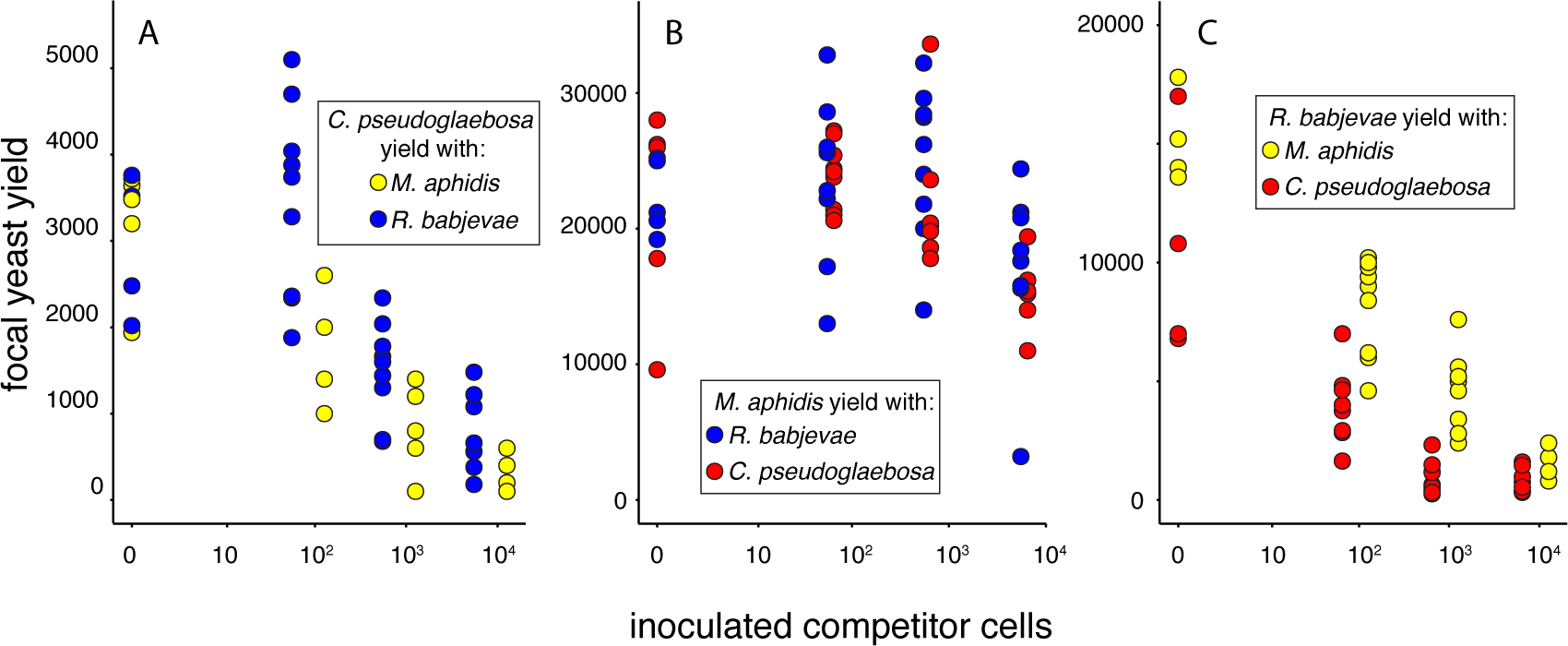
Influence of interacting species on (A) *C. pseudoglaebosa*, (B) *M. aphidis*, and (C) *R. babjevae*. The plots depict the yield of each focal species as a function of the number of cells of an interacting species co-inoculated with the focal species. Interacting species are coded by color: red = *C. pseudoglaebosa*, yellow = *M. aphidis*, and blue = *R. babjevae*.

### C. pseudoglaebosa *is an early disperser in pitchers*

To investigate whether dispersal influences *C. pseudoglaebosa* dominance in pitchers, we observed the arrival times each of the three yeasts mentioned above in pitchers over the *S. purpurea* growth season in Harvard Pond. We surveyed the presence or absence of each yeast in each of the 43 sampled pitchers using taxon-specific PCR primers (Table 1) to determine when each yeast arrived in a pitcher and whether it persisted throughout the season. The three yeasts appeared in pitchers sequentially (Figure 6): *C. pseudoglaebosa* first arrived in pitchers within four days after the pitchers opened; *R. babjevae* arrived between four days and one week after pitchers opened; and *M. aphidis* arrived one week to one month after pitchers opened. Once a yeast colonized a pitcher, it either persisted in or disappeared from that pitcher later in the season, but it did not disappear from the broader metacommunity.

**Figure 6:**
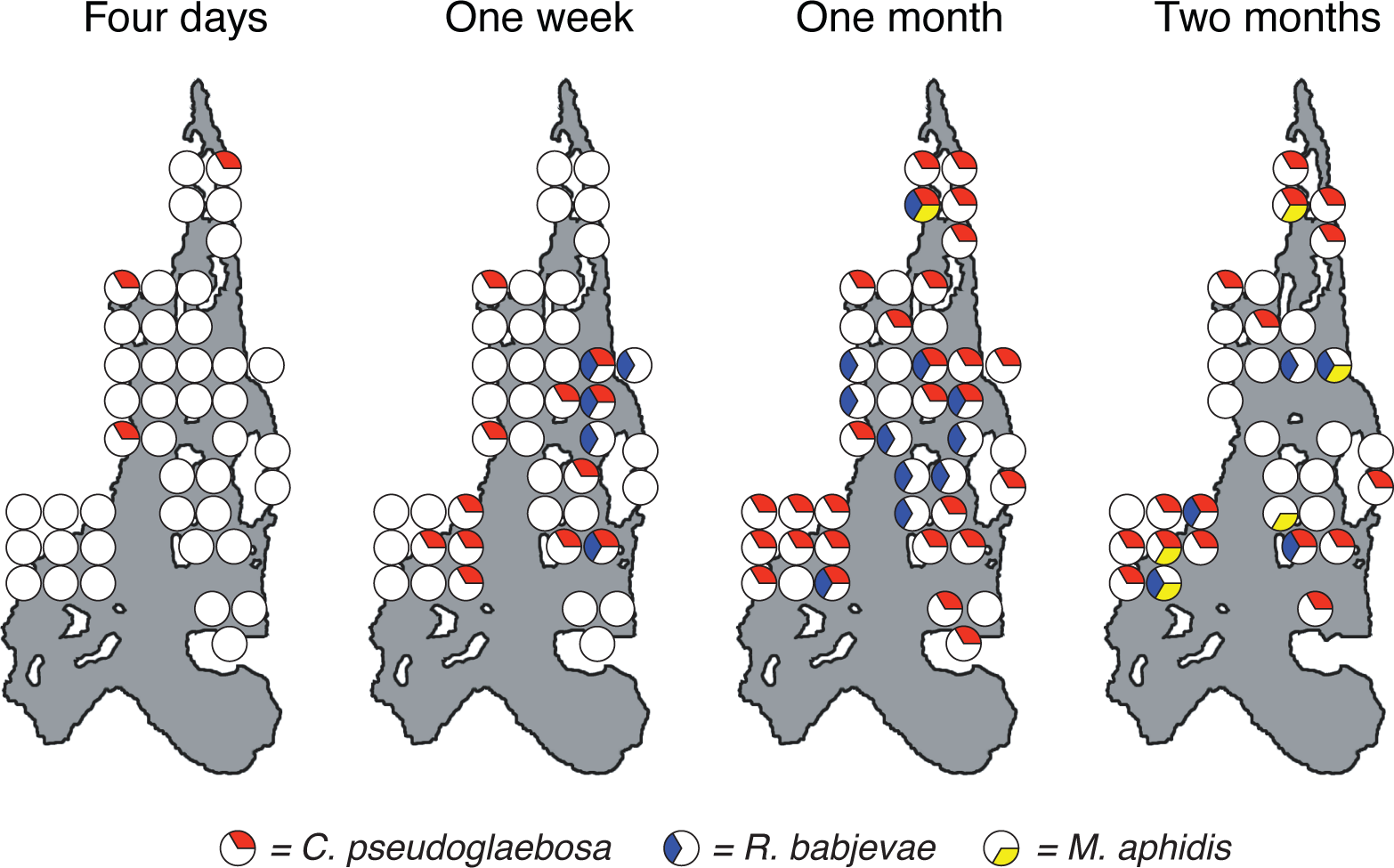
Presences and absences of each of three yeasts in 43 pitchers over time. Each large gray shape represents the Harvard Pond at one of four time points. Circles represent pitchers: completely white circles represent pitchers in which none of the three yeasts were detected, and circles containing colored pie slices represent pitchers in which one or more of the three assayed yeasts were detected. Pie slices are colored by detected yeast: red = *C. pseudoglaebosa*, yellow = *M. aphidis*, and blue = *R. babjevae*. Circles indicate the approximate locations of pitchers, and are offset to make all data visible; see Figure 1C for accurate pitcher locations.

## Discussion

### C. pseudoglaebosa *is a pitcher metacommunity dominant*

*C. pseudoglaebosa* was the dominant fungal taxon throughout succession in the *S. purpurea* pitcher metacommunity, although it did not dominate every individual pitcher community. It was the most frequent taxon in the metacommunity at every sampled timepoint, and its frequency in the metacommunity increased after initial colonization (Figure 4). Dominance in the pitcher metacommunity was in part a result of the metacommunity structure itself: while *C. pseudoglaebosa* was not dominant in every pitcher, it was dominant in enough pitchers (in some cases with a frequency above 90% of all OTUs) to dominate the metacommunity as a whole (Figure 4). The poor competitive performance of *C. pseudoglaebosa* relative to other yeasts in microcosms suggests that overall competitive superiority was not the cause of its dominance (Figure 5). Instead, *C. pseudoglaebosa*’s early dispersal is a more likely cause (Figure 6).

Our best explanation for *C. pseudoglaebosa*’s dominance throughout succession in the pitcher metacommunity is that it pre-empted other community members by dispersing and establishing in pitchers before other taxa. Early dispersal likely gave *C. pseudoglaebosa* a numerical advantage by providing the opportunity to begin exponential growth before other members of the fungal community could arrive and begin growing (Scheffer *et al.* 2017). Additionally, once *C. pseudoglaebosa* was established in a pitcher, facilitation by low-frequency interacting taxa may have helped it maintain dominance (Figure 5A). Once established early in the metacommunity, *C. pseudoglaebosa* continued to disperse into new pitchers from one week to one month into the approximately two month long growing season (Figure 6).

Although *C. pseudoglaebosa* was the numerically dominant fungal taxon in the metacommunity of pitchers, chance events, dispersal, and interactions among fungi determined whether it was the dominant taxon inside any given pitcher. We expect that interactions with taxa that arrive, by chance, at different times in different pitchers caused the variety of *C. pseudoglaebosa* relative frequency changes observed (*i.e.,* increasing, decreasing, or nonmonotonic, Figure 4B) because interacting yeasts have qualitatively different impacts on *C. pseudoglaebosa* performance depending on the number of interacting cells present (Figure 5A). Individual pitchers experienced priority effects because the timing of taxon arrival in each pitcher (*e.g.,* early arrival of *C. pseudoglaebosa* in a pitcher) determined later community composition (*e.g., C. pseudoglaebosa* dominance in the pitcher) (Fukami 2015). Despite the influence of chance events on *C. pseudoglaebosa* dominance in any given pitcher, the early and frequent dispersal of *C. pseudoglaebosa* compared to other yeasts enabled its overall dominance in the metacommunity (Figure 4A, 6).

### Ecological patterns and processes during pitcher metacommunity succession

In the metacommunity, and in many individual pitchers, *C. pseudoglaebosa* remained dominant through decreases in fungal taxon richness and diversity (Figures 3, 4). We did not observe a hump-shaped relationship between pitcher age and species richness, as previously predicted (Figure 3A) (Moquet *et al.* 2003). Species richness instead decreased, even as *R. babjevae* and *M. aphidis* were first dispersing into pitchers late in the season (Figures 3A, 6). It is likely that *C. pseudoglaebosa* repression of taxa through priority effects has a larger influence on species richness than increasing diversity through new dispersal does.

Previous studies have also documented biotic and abiotic successional changes in pitchers; while we did not measure the same parameters as these previous studies, we assume that similar changes occurred in our pitcher metacommunity and that *C. pseudoglaebosa* maintained its dominance through these changes. For example, previous studies documented decreasing pH with increasing pitcher age, an early peak in prey insect capture during pitchers’ life spans (Fish & Hall 1978), and a variety of changes in bacterial, protist, and invertebrate community compositions over time (Gray *et al.* 2012, Miller & terHorst 2012). In bacterial, protist, and invertebrate communities, the identities of dominant taxa changed as succession progressed. In contrast, *C. pseudoglaebosa* remained the dominant fungus throughout succession. *C. pseudoglaebosa* appears to be a classical early-successional taxon (Connell & Slatyer 1977, Fierer *et al.* 2010) because it disperses early and frequently (Figure 6), but it is not replaced by late-successional taxa.

*C. pseudoglaebosa* dominance throughout succession may be enabled by the short lifespans of pitchers in Harvard Pond; *i.e.,* pitchers may not live long enough to enable late-succession fungal taxa to dominate the metacommunity. We sampled pitchers that were up to 66-74 days old, and stopped sampling at this age because 23% of pitchers had been destroyed by moths. However, pitchers in northern *S. purpurea* populations can survive intact through winter conditions (Judd 1959), and pitchers can be active for over a year in the southern United States (Miller & terHorst 2012). We speculate that fungal succession more closely resembles classical successional patterns and the patterns observed for other pitcher guilds (*e.g.,* bacteria, invertebrates) in longer-lived pitchers. For example, it is possible that a strong competitor such as *M. aphidis* could replace *C. pseudoglaebosa* in southern *S. purpurea* metacommunities where pitchers are active for many months. However, consistent dominance of a single taxon over succession may be common in microbial habitats that, like northern *S. purpurea* pitchers, have short lifespans but repeatedly become available.

## Conclusions

In the model pitcher plant metacommunity, taxon dispersal ability has a profound influence on community structure. In particular, *C. pseudoglaebosa*’s ability to disperse into pitchers before other fungal taxa enables it to persist as the dominant taxon in the pitcher metacommunity, even as intertaxon interactions and the stochasticity of individual dispersal events prevent its dominance in every pitcher. It is likely that dispersal ability leads to persistent dominance in a variety of other natural succeeding microbial communities and metacommunities, especially when early dispersal allows a taxon to prevent establishment of other taxa.

Future studies of microbial succession should explicitly include metacommunity structure when investigating ecological processes. In the pitcher metacommunity, overall taxon composition changed little over time, with *C. pseudoglaebosa* dominant throughout succession (Figure 4A). However, individual pitchers followed a variety of trajectories (Figures 2, 4B, 6). Studies of succession that do not take a metacommunity’s structure into account may miss community heterogeneity and the diversity of ecological processes, especially dispersal ability, in play among communities.

## Materials and Methods

### Study site and field collections

Observations were made on *Sphagnum* islands in Harvard Pond, adjacent to Tom Swamp, a 50 ha *Sphagnum* bog located in Petersham, Massachusetts at 42°30’N, 72°12’W (Figure 1; Swan & Gill 2007). The *C. pseudoglaebosa* and *M. aphidis* isolates used in the microcosm study were collected from pitchers in Harvard Pond, and the *R. babjevae* isolate was collected from a pitcher in Swift River Bog, a 2 ha kettlehole bog located 75 km south of Tom Swamp in Belchertown, MA at 42°16’N, 72°20’W (Ellison *et al.* 2002). These three yeast isolates were collected in the summer of 2006 and identified by comparing their ribosomal sequences, amplified using the ITS1F/ITS4 and LS1/LR5 primer pairs (Gardes & Bruns 1993, White *et al.* 1990, Hausner *et al.* 1993 Vilgalys & Hester 1990), to sequences in the NCBI BLAST database (Zhang *et al.* 2000). We chose *C. pseudoglaebosa*, *R. babjevae*, and *M. aphidis* in part because they were all easily cultured from pitchers, and in part because they formed colonies with different morphologies on agar plates: *C. pseudoglaebosa* forms smooth white colonies; *M. aphidis* forms wavy white colonies; and *R. babjevae* forms smooth pink colonies.

All *S. purpurea* pitcher water samples for PCR and 454 sequencing were collected in the spring and summer of 2009. In May of 2009, we identified 43 unopened *S. purpurea* pitchers on 32 *Sphagnum* islands in Harvard Pond. Pitchers ranged from less than 1 m to 908 m in distance to other pitchers (Figure 1C). We visited each pitcher daily until it opened, and counted pitcher age from the date it opened. For each pitcher water collection, the water inside a pitcher was first mixed by pipetting up and down with a sterile plastic transfer pipette. We then removed about 0.25 ml pitcher water and mixed it with 0.25 ml of CTAB buffer (100 mM Tris pH 8.0, 1.4 M sodium chloride, 20 mM EDTA, 2% CTAB). To the best of our ability, we avoided collecting insect prey or macrofauna in these samples, although any protists and microscopic animals present in our samples were included; collected pitcher water contained no large animal parts and appeared as a cloudy liquid. All samples were flash-frozen in liquid nitrogen within five hours of collection and stored at −20 or −80°C before DNA extraction.

### PCR assay

We assayed each pitcher water sample for amplifiable DNA from all fungi, using the ITS1F/ITS4 primer pair, and for each of the three yeasts in the microcosm experiment, using the primers in Table 1. Primers to selectively amplify portions of each microcosm yeast’s ITS sequence were designed using the NCBI BLAST primer tool (Rozen & Skaletzky 2000). We chose primer sequences to reliably amplify as much of the ITS sequence of each yeast species as possible, while not amplifying other sequences in the BLAST database.

To extract DNA from each pitcher water sample before the PCR assay, we first thawed and centrifuged frozen samples at 16.1 g for 10 min and removed the supernatant from each pellet. We then suspended each pellet in 200 µL of breaking buffer (2% Triton X-100, 1% sodium dodecyl sulfate, 100 mM sodium chloride, 10 mM Tris, and 1 mM EDTA) (Hoffman 1997). We mixed each suspension with about 200 µL of 0.5 mm glass beads and 200 µL 25:24:1 chloroform:phenol:isoamyl alcohol. We vortexed each mixture for 2 min, and then centrifuged it for 5 min at 16.1 g. After centrifugation, we removed the aqueous layer and mixed it with 2.5 volumes of 95% ethanol and 0.1 volume of 3M sodium acetate (Sambrook & Russell 2001); we incubated each aqueous layer mixture at −20°C for at least three hours. Next, we centrifuged each aqueous layer mixture for 15 min at 16.1 g, and removed the supernatant. Finally, we washed each pellet with 0.5 ml 70% ethanol, centrifuged each mixture for 10 min at 16.1 g, removed the supernatant, and resuspended each pellet in 50 µl water.

We then assayed each DNA extract for the presence of each Fungal taxon, or any Fungal DNA in the case of the ITS1F/ITS4 primer pair, using PCR. Each PCR reaction was composed of 7.9 µL water, 0.1 µL GoTaq^®^ Flexi polymerase (Promega), 5 µL Flexi buffer with green dye added, 5 µL 5x CES (combinatorial PCR enhancer solution: 2.7 M betaine, 6.7 mM dithiothreitol, 6.7% dimethyl sulfoxide, 55 µg/mL bovine serum albumin) (Ralser *et al.* 2006), 5 µL nucleotide mix, 2 µL magnesium chloride, 1 µL of 10 µM of each primer, and 1 µL undiluted template DNA extract. All reactions were cycled on a Biorad iCycler or myCycler using denaturing, annealing, and extension temperatures of 95, 55, and 72 °C, respectively. We denatured for 85 s, then ran 13 cycles of 35 s denaturing, 55 s annealing, and 45 s extension, followed by 13 cycles that were identical but had a 2 min extension, and finally 9 cycles with a 3 min extension. We ran a subsequent 10 min extension. Two µL of each PCR product were visualized on 1% agarose gels stained with SYBR Safe dye (Invitrogen) and photographed using a U:genius gel documenting system (Syngene) and a Stratagene transilluminator. Photographs of gels were scored for presence or absence of a band. Bands that were too faint to reliably score were run a second time with 6 µL of PCR product per well. Presence of a band on a gel indicated the presence of detectable fungal or yeast species DNA in a water sample.

To confirm that primers only amplified sequences from the target yeasts, we randomly selected nine PCR products generated from the *C. pseudoglaebosa* and *R. babjevae* primer pairs for sequencing. The primer pair that targets *M. aphidis* only amplified DNA from six pitcher water extracts, and we sequenced all six PCR products for this primer pair. Sequences were identical to or within one base of cultured isolate sequences.

### Pitcher water fungal DNA amplification and 454 sequencing

We extracted and amplified fungal DNA for fungal community amplicon sequencing using the protocols described above, with the following changes. Gotaq^®^ Hotstart polymerase (Promega) was used instead of Flexi polymerase, and we used 50 µM instead of 10 µM of the reverse primer. The forward primer consisted of (in order from 5’ to 3’) the 454 “A” primer (CCATCTCATCCCTGCGTGTCTCCGACTCAG) concatenated with a 10-bp multiplex tag (454 Life Sciences Corporation 2009), and ITS4; the reverse primer consisted of the 454 “B” primer (CCTATCCCCTGTGTGCCTTGGCAGTCTCAG) concatenated with ITS1F. Multiplex tags were unique to each sample. Reactions were cycled at 95 °C for 15 min; 30 cycles of 95 °C for 1 min, 51 °C for 1 min, 72 °C for 1 min; and a final extension of 72 °C for 8 min.

Products were purified using Agencourt^®^ *AMPure*^®^ XP (Beckman Coulter) and quantitated using a Qubit^®^ dsDNA HS Assay (Invitrogen) according to the manufacturers’ instructions. We combined equimolar concentrations of the products of each of three separate PCR reactions from each DNA extract. The sequencing pool consisted of pooled equimolar concentrations of each pooled PCR product. The pool was sequenced on one-eighth of a 454 Titanium sequencing run by the Duke Genome Sequencing & Analysis Core Resource.

### 454 sequence processing

We processed sequences using QIIME 1.3.0 (Caporaso *et al.* 2010). Low quality sequences were removed and the remaining sequences were assigned to mulitplex barcodes using the default quality filtering settings. Primers and barcodes were trimmed from each sequence and sequences shorter than 200 and longer than 1000 bp were removed from each dataset. Sequences were denoised using the QIIME denoiser. We reduced chimeric sequences by trimming the 5.8s and ITS1 portions from all sequences using the Fungal ITS Extractor (Nilsson *et al.* 2010a), and only analyzing the ITS2 portion. The 5.8s ribosomal region lies between the ITS1 and ITS2 spacers, and is conserved among fungi relative to the spacers. We expected most chimeric sequences to form in the 5.8s region and to be composed of ITS1 and ITS2 sequences from different templates (Nilsson *et al.* 2010b). We chose operational taxonomic units (OTUs) using the uclust method in QIIME, at 97% similarity (Edgar 2010). We discarded all OTUs composed of a single sequence (singleton OTUs) because we assumed that they resulted from sequencing errors. The longest sequence in each remaining cluster was retained as a representative sequence. Of the 141,424 total sequences produced, 27,632 were discarded for having lengths less than 200 or more than 1000 bp and 10,938 were discarded because they had low quality or did not have a matching barcode. Sixty-six sequences were discarded because the ITS2 subunit could not be extracted and 139 sequences were discarded because they represented singleton OTUs. In total, we retained 102,649 sequences for further analysis. Each pitcher water sample produced between 253 and 4365 sequences. We will deposit the 454 sff files and corresponding FASTA files in the NCBI Small Read Archive database (accession numbers to be added after manuscript acceptance).

### Sequence taxonomy assignments

To assign genus-level taxonomy to OTUs, we performed a MEGAN analysis of OTU representative sequences on the top ten hits from the NCBI BLAST nucleotide database extracted using BLAST 2.2.25+ and default MEGAN settings (Huson *et al*. 2011, Zhang *et al.* 2000). We discarded OTUs matching organisms not in the kingdom Fungi (plant, animal, and protist sequences), but we assumed that OTUs with no BLAST matches or matching unassigned fungal environmental sequences were fungal sequences not yet identified in the NCBI database. We retained unassigned OTUs for diversity measurements, but did not include them in taxonomy summaries. We reviewed the taxon assignments output by MEGAN manually, and filled in higher-level classifications (*e.g.,* order or class) using Index Fungorum (http://www.indexfungorum.org/) when an OTU was assigned to a genus but not higher-level classifications. We attempted to assign all taxa to genera; if it was not possible to assign a taxon to a genus, we assigned it to the most specific taxonomic group possible. OTUs assigned to the genus *Candida* were further assigned to the species *C. pseudoglaebosa* if they matched either *C. pseudoglaebosa* or its close relative *C. glaebosa* in a BLAST search. When we compared these seven OTUs against sequences representing the type strains of the two *Candida* species (accession numbers KY102342.1 and KY102112.1, respectively), all seven OTUs aligned to *C. pseudoglaebosa* better than they did *C. glaebosa*. In total, we detected 553 OTUs, of which 379 were assigned to fungal taxa; fifteen were discarded because they matched non-fungal sequences; and 159 were not assigned. Of the 379 fungal taxa in the temporal data set, 52% were Basidiomycota, 43% Ascomycota, and 5% basal fungal lineages. Sequences, metadata, and OTU tables including taxonomic assignments in QIIME 1.3.0 format are included in the supplementary datasets 2-3.

### Microcosm interaction assays

Interactions between yeasts were assayed in microcosms designed to mimic pitchers simultaneously colonized by different numbers of two yeast species. We grew microcosms in low-nutrient media designed to mimic natural conditions in pitchers. Microcosms contained sterile yeast extract media (YEM) composed of 1g/L yeast extract in local tap water (Cambridge, MA, USA). Tap water was used instead of deionized water because we wanted the media to include micronutrients present in local rainwater that may be important for pitcher plant yeast growth. The tap water supply in Cambridge, MA, where this experiment was conducted, comes from three Massachusetts reservoirs (Waldron & Bent 2001), and we expected it to have similar inputs as rainwater in Harvard Pond pitchers. Each microcosm was inoculated with a target yeast species and an interactor in 200 µL of liquid yeast media. Each target yeast was inoculated with about 1000 cells per microcosm, and each interactor yeast was inoculated at zero, low, medium, and high cell numbers (0 and approximately 100, 1000, and 10,000 cells).

Eighteen treatments of yeast mixtures were prepared, with ten replicates each, for a total of 180 microcosms. Before inoculation, yeasts were grown in liquid YEM for 48 hours. Inoculation sizes were measured after inoculation using counts of colony-forming units (CFUs) on solid YEM (YEM plus 1.5% agar). Microcosms were arranged in sterile 96-well polystyrene flat bottom cell culture plates and incubated between 25 and 27 °C, shaking at 700 rpm for 48 hours. After incubation 32 microcosms were discarded because of suspected contamination. We diluted each remaining microcosm 1:10^3^ or 1:10^4^ in sterile water, plated it to solid YEM, and counted CFUs on plates containing at least 30 total CFUs. When no CFUs of an inoculated yeast were present on a plate, we conservatively assumed that the yeast was present in the microcosm in numbers just below our detection limit. We calculated total cell numbers assuming one instead of zero CFUs for these yeasts absent from plates. CFU counts before and after incubation are included in supplementary dataset 4.

### Statistical analyses

OTU datasets rarefied to 1140 sequences were used to produce Non-metric Multidimensional Scaling (NMDS) plots and to compare community similarities, and alpha diversity indices among pitchers. Eight samples contained fewer than 1140 sequences and were discarded. Proportions of samples assigned to taxonomic groups were calculated based on the full non-rarefied dataset. Community similarities over time were compared using partial distance-based redundancy analysis (db-RDA) of Jaccard dissimilarity (Jaccard 1912) between each pair of samples with pitcher age as the explanatory variable, conditioned on pitcher identity. A correlation between geographic distance and community similarity was conducted using a partial Mantel test conditioned on pitcher age. Hill numbers of order *q*=0 or 2 (^q^D) were calculated as 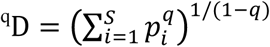, where *S* is the total number of OTUs and *p*_i_ is the relative abundance of OTU *i*; ^1^D was calculated as the exponent of Shannon diversity (Hill 1973, Chao *et al.* 2014). Changes in Hill numbers were modeled over time using repeated-measures linear models controlled for pitcher identity; ^1^D and ^2^D were log-transformed before analyses to homogenize variances among timepoints, and ^0^D was not transformed.

We modeled the impact of interactor yeasts on focal yeasts in microcosms using multiple linear and polynomial regressions. Separate regressions were conducted for each focal yeast. For each regression, focal yeast yield was the dependent variable, and both the number of co-inoculated interactor yeast cells and the identity of the interactor yeast were independent variables. We modeled both linear and quadratic relationships between the number of co-inoculated interactor yeast cells and the dependent variable because the relationship did not always appear linear when plotted. Before constructing the regressions, we square-root-transformed focal yeast yield to homogenize variances for the focal yeasts *R. babjevae* and *C. pseudoglaebosa*, but left yield untransformed for the focal yeast *M. aphidis*. We also transformed competitor inoculum size by log_10_(x+1) because interactor inoculum size was varied on a log scale in the experiment. When comparing the influences of competitor species, we randomly assigned treatments with no interacting yeast inoculum to one of the two interacting yeasts. When selecting the best-fitting regression model, we first established the best-fitting relationship (linear, quadratic, or both) between log-transformed interactor inoculum size and focal yeast yield, and then determined whether adding interactor identity or interactions between interacting yeast identity and inoculum size to the model improved it. The best-fitting model was the one with the lowest Akaike Information Criterion (AIC).

All statistical analyses and index calculations were conducted using R version 3.3.1 (R Development Core Team, 2016) and the packages vegan, fields, nlme, and GUniFrac (Chen 2018, Pinheiro *et al.* 2016, Oksanen *et al.* 2016, Nychka *et al.* 2016) plots were made using ggplot2 (Wickham 2016)

## Acknowledgments

Thank you to Aaron Ellison for help throughout this project, and especially for helpful comments on this manuscript. We would also like to thank staff and REUs of Harvard Forest for help in the field; members of the Marx lab for advice on analyzing competition data; Chris Hittinger for advice on *Candida* taxonomy; and Tadashi Fukami, and the Pringle lab for helpful comments on the manuscript. We would like to thank Harvard Forest and the Massachusetts Department of Fish and Wildlife for fieldwork permissions. Funding was provided by Harvard University and a National Science Foundation Doctoral Dissertation Improvement Grant (award ID number 0909694).

---

Figure 4 Figure Supplement 1: Taxon diversity in pitchers over time; rarefied sequence data. The OTU table was rarefied to 1140 sequences per sample. (A) Taxon diversity in the entire bog metacommunity. Colored bars represent proportions of total sequences for each fungal class (or phylum for basal fungal lineages). The hatched area represents total *C. pseudoglaebosa* frequency for each time point. Note that *C. pseudoglaebosa* is in the class Saccharomycetes and represents over 99% of Saccharomycetes sequences at the one and two month time points. (B) *C. pseudoglaebosa* sequence frequency in individual pitcher communities. Data points for communities in the same pitcher are connected with lines. Black lines connect points for pitcher group 1 and gray lines connect points for pitcher group 2. Note that data for some pitchers are missing at some time points because these samples were discarded during sample rarefaction.

